# Two-way migration of *Lychnis wilfordii* caused by the circular landform of Japan-Korea-northeast China-Russian Far East region and its suggestion for conservation in northeast Asia

**DOI:** 10.1101/857227

**Authors:** Saya Tamura, Myounghai Kwak, Goro Kokubugata, Chan-ho Park, Byoung-Yoon Lee, Tomoko Fukuda, Ekaterina Petrunenko, Inna Koksheeva, Elena Pimenova, Pavel Krestov, Svetlana Bondarchuk, Jin-Shuang Ma, Hai-Cheng Zhou, Hayato Tsuboi, Yoko Nishikawa, Takashi Shimamura, Hiroko Fujita, Koh Nakamura

## Abstract

In northeast Asia, substantial portion of the floras, including endangered species, are shared among its component countries in the continental, peninsula, and island parts largely through Quaternary migration. To effectively conserve nationally endangered plants in Northeast Asia, transnational conservation studies are vitally needed. *Lychnis wilfordii* (Caryophyllaceae) has disjunct distribution in Russian Far East (Primorsky Krai), northeast China (Jilin), Korea (Gangwon-do) and Japan (Hokkaido, Aomori, Nagano), surrounding the sea, and this is designated as an endangered species in Japan and Korea. Population genetic and molecular dating analyses were conducted 1) to elucidate geographic genetic structure covering the species range, 2) to test possible scenarios of migration, and 3) to develop logical plans for effective conservation. Population genetic analyses indicated the continent and peninsula parts (north and south Primorsky Krai, Jilin, and Gangwon-do) had higher genetic diversity compared to those in the Japanese Archipelago (Hokkaido and Nagano). Five genetically distinct groups were recognized, namely, Nagano, Gangwon-do, Jilin, north and south Primorsky Krai plus Aomori, and Hokkaido. Genetic distance between Hokkaido and Nagano was larger than between Hokkaido and north Primorsky Krai, and between Nagano and Gangwon-do, crossing national borders and the natural barrier of the sea. Considering these results, *L. wilfordii* likely migrated from the Asian continent to the Japanese Archipelago using two routes: north route from Russian Far East to Hokkaido and Aomori, and south route from the Korean Peninsula to Nagano. Based on molecular dating, migration from the continent to the islands likely occurred from the middle Pleistocene to the Holocene. For effective conservation of *L. wilfordii*, Hokkaido and Nagano populations should be distinguished as different evolutionary significant units, although these two regions belong to the same country, because Hokkaido and Nagano populations are at the different ends of the two migratory routes based on the migration scenario.

## INTRODUCTION

Flora of northeast Asia, here defined as Japan-Korea-northeast China-Russian Far East region, is rich and occupies a unique position in global plant diversity, containing many relic and neo-endemic plant lineages (Good, 1974; Maekawa, 1974; Takhtajan, 1986; Wu & Wu, 1996; Tang & Ohsawa, 2002). At the same time, considerable portions of the floras of the component countries (ca. 10-25% in each country) are currently threatened or on the verge of extinction due to extrinsic factors (habitat loss and degradation, overexploitation, climate change) as well as intrinsic ones (limited habitat due to unique ecological demands) (Biodiversity working group of China council for international cooperation on environment and development, 2004; Community of natural resources and environment conservation of Sakhalin state *et al.,* 2005; Administration of Kamchatka region *et al.,* 2007; Administration of Primorsky Krai krai *et al.*, 2008; National institute of biological resources, 2014; Ministry of the Environment, Government of Japan, 2015). These countries share substantial portion of the floras (Good, 1974; Takhtajan, 1986), and plants designated as endangered species in one country are not all national endemics but distributed in neighboring countries as endangered or non-endangered species (Kokubugata, 2016). Nevertheless, conservation studies on endangered plants in this region are usually confined to one country (Jin *et al.,* 2003; Washitani *et al.,* 2005; Hu *et al.,* 2010; Jae & Sungwon, 2016) and the progress of plant conservation in northeast Asia is impeded by national borders. Rare examples of pan-northeast Asian studies include an attempt to compile an integrated red list of East Asian plants, including Russian Far East (Kokubugata, 2016). For endangered plants distributed across national borders, conservation of populations in one country but not in others is ineffective if those populations share the same gene pool because this would lead to genetic diversity loss in “conserved” populations once nonconserved ones are lost. To effectively conserve nationally endangered plants in northeast Asia, that often have conspecific populations abroad, transnational conservation studies are vitally needed.

A unique feature of endangered plant species in northeast Asia is its disjunct distribution in the continental part (northeast China and Russian Far East), the Korean peninsula, and the islands (in particular, the Japanese Archipelago and Sakhalin), encircling a geographic barrier of the sea. Geohistorically, due to Quaternary sea level fluctuations, the Japanese Archipelago was repeatedly connected to the Eurasian continent with landbridges via Sakhalin until the Last Glacial Maximum in the north (Pietsch *et al.*, 2003) and via the Korean peninsula until the late Pleistocene in the south (Ohsima, 1990), forming a continuous circular landform. Many plant species migrated from the continent to the Japanese Archipelago via these landbridges during Quaternary, as revealed by molecular studies (e.g. alpine plants: Fujii & Senni, 2006; Ikeda *et al.*, 2008, trees and shrubs: Okaura *et al.*, 2007; Aizawa *et al.*, 2007, 2009, herbs: Lihová, *et al.*, 2010; Kikuchi *et al.* 2010, 2013). These studied plants are mostly common species, and few studies focused on endangered plants for the conservation. Elucidating genetic legacy from migration/range expansion in the landform of northeast Asia can contribute substantially to increase the effectiveness of conservation planning based on genetic profile of populations because it is not only the result of population-genetic processes in ecological time scale. Broad spatial and temporal scale studies are needed for endangered plants in northeast Asia.

*Lychnis wilfordii* (Regel) Maxim. (Caryophyllaceae) is one of such endangered species that has disjunct distribution in northeast Asia surrounding the sea, i.e. in Russian Far East (Primorsky Krai), northeast China (Jilin), Korea (Gangwon-do) and Japan (Hokkaido, Aomori, Nagano) (Figs. 1, 2; Lu *et al.*, 2001; Aomori prefecture, 2010; National Institute of Biological Resources, 2014; Tamura *et al.,* 2016; Kadota, 2017). This species grows in wet meadows and at wet woodland edges. The numbers of individuals and habitats of this species are decreasing due to ecological succession, exploitations, and overcollection for horticultural purposes, and this species is designated as an endangered species in Japan and Korea (National Institute of Biological Resources, 2014; Ministry of the Environment, Government of Japan, 2017). In northeast China, it is not designated as an endangered species but treated as a rare species. In Russian Far East, on the other hand, this is a common species and there are larger number of individuals and populations.

**Figure 1.**
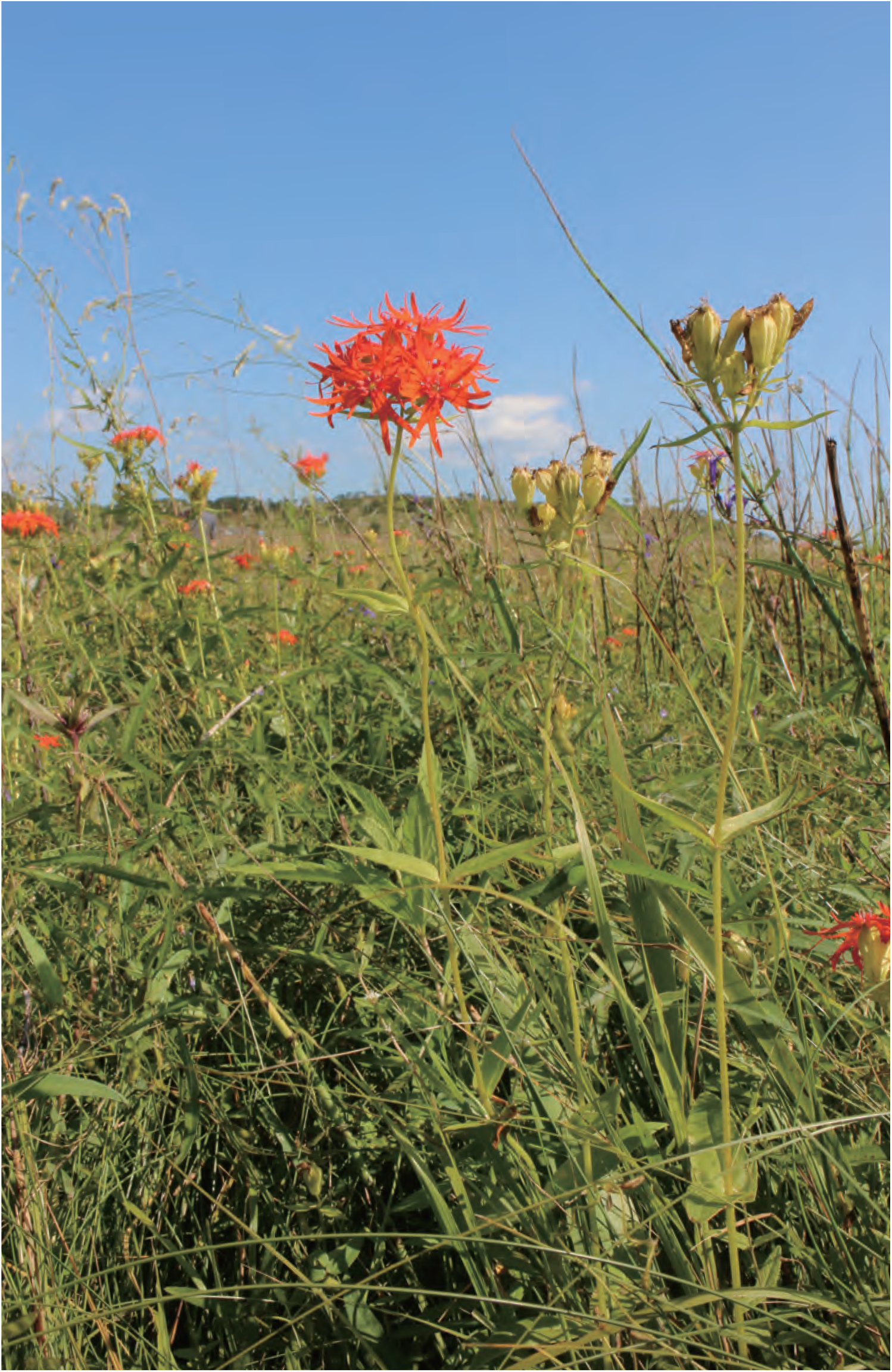
*Lychnis wilfordii* in a habitat in Russia Far East (Sep. 3, 2017).

**Figure 2.**
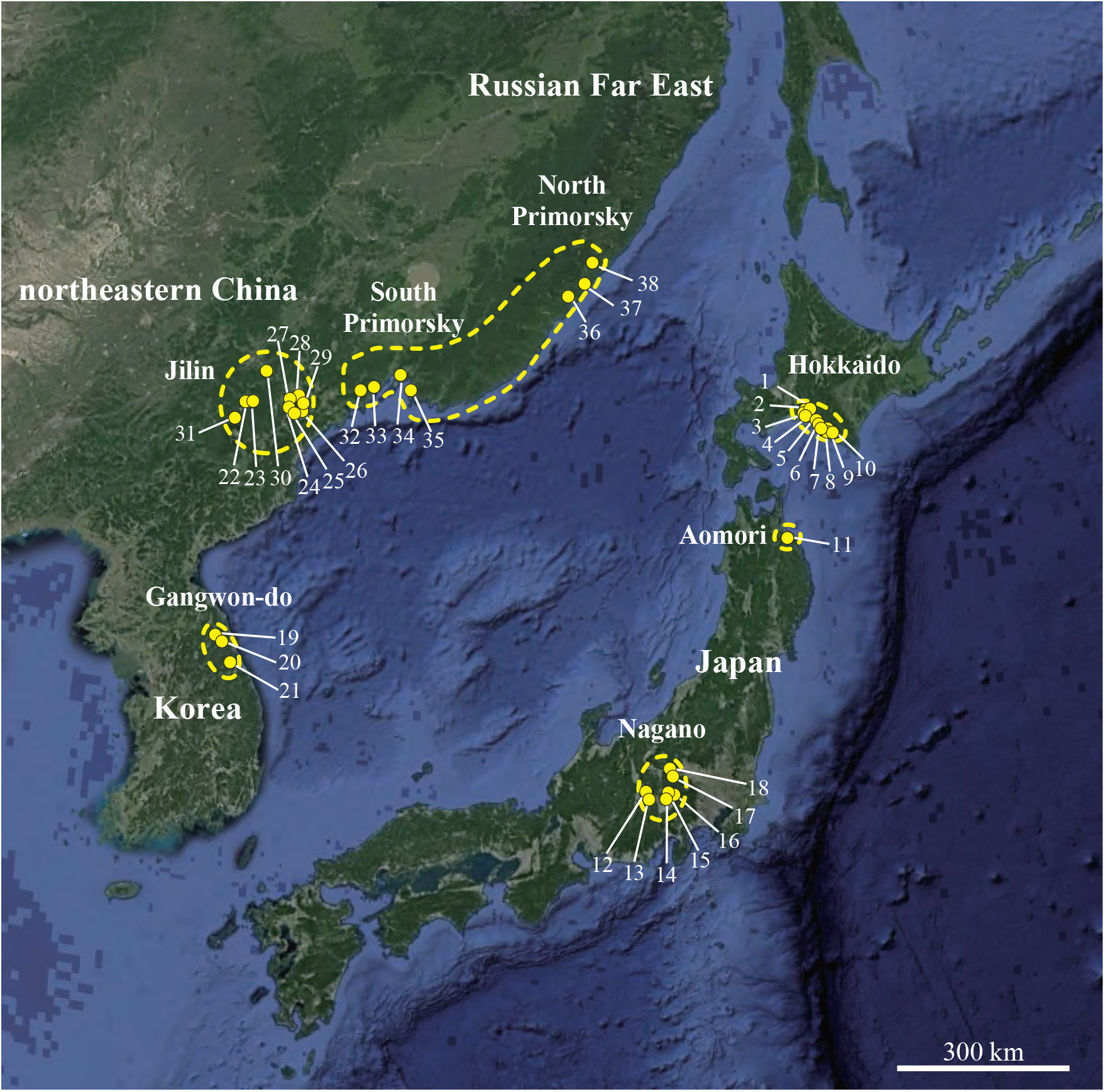
Species range and collection sites of *Lychnis wilfordii*. Circles of broken line show the species range and small filled circles show the sampling populations. For the population numbers (1-38), see Table 1.

**Table 1.**
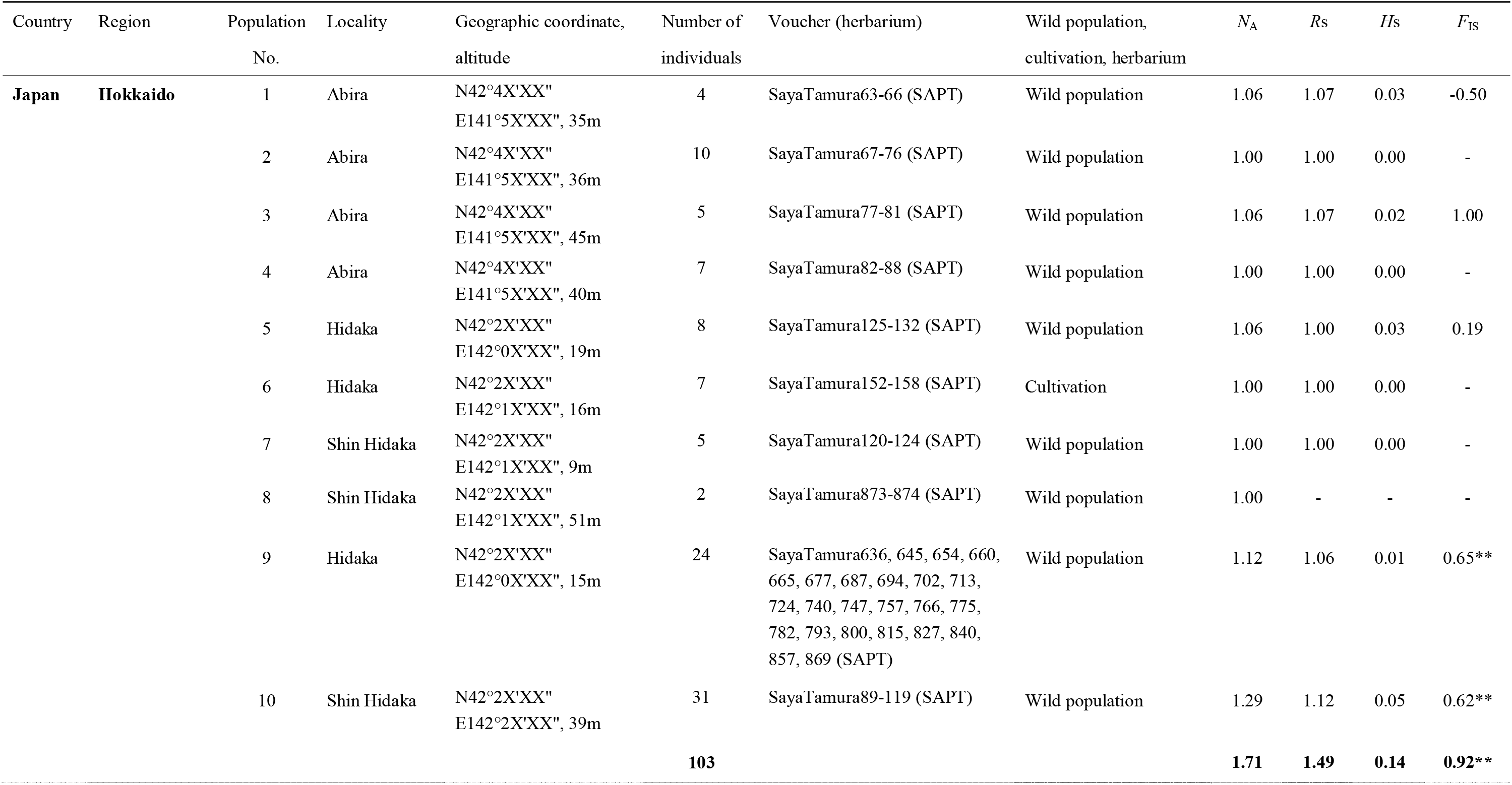
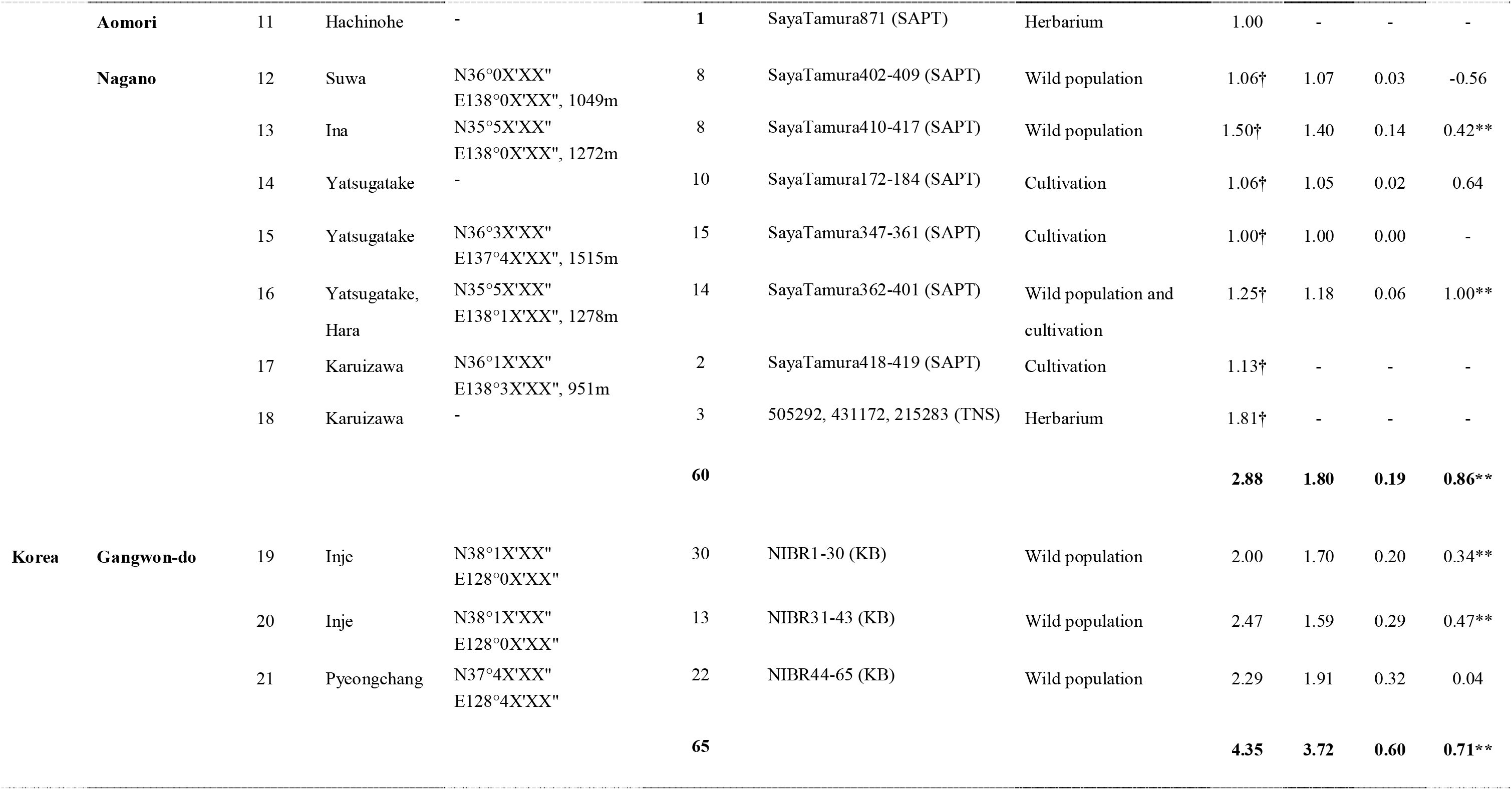
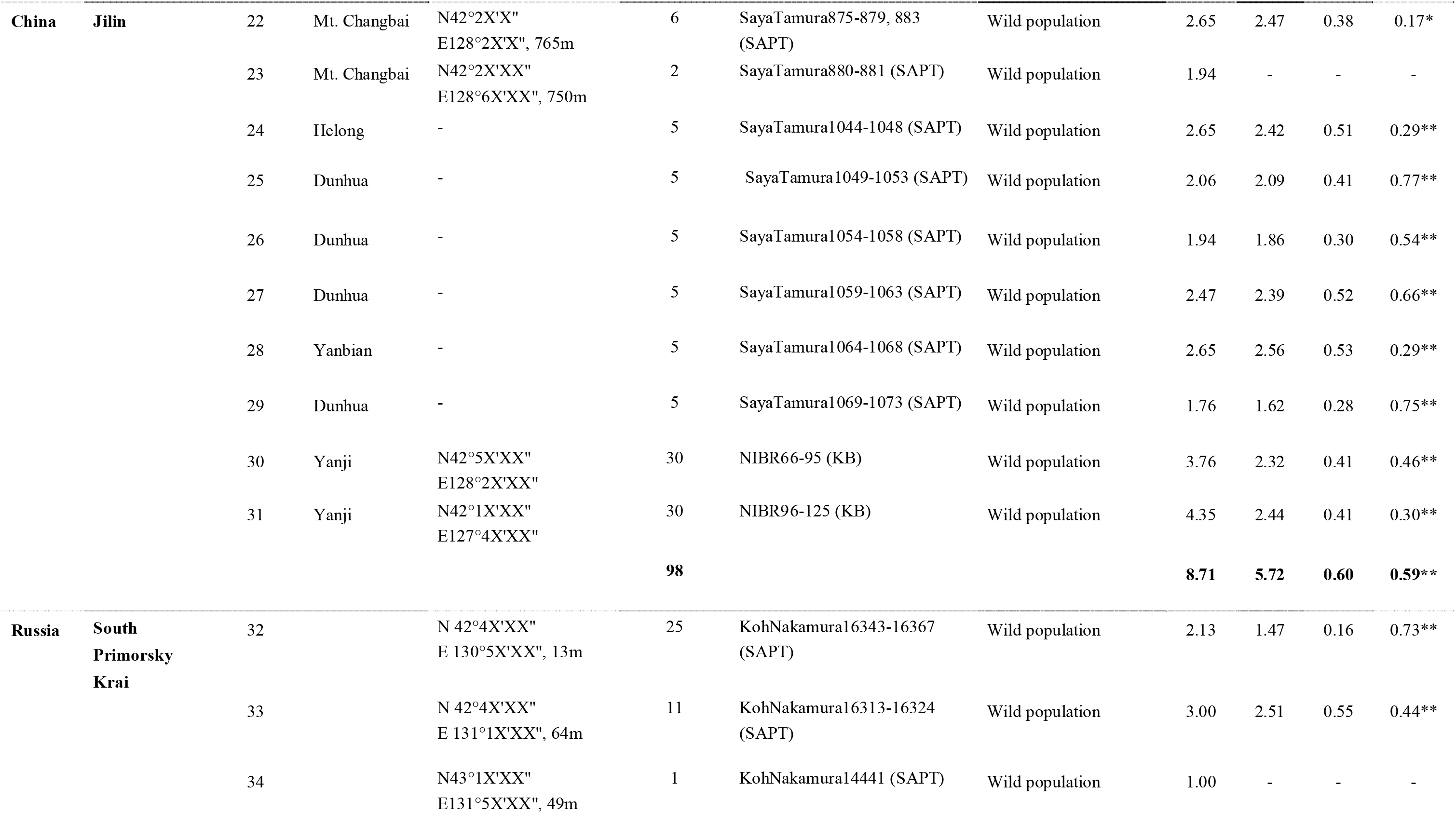
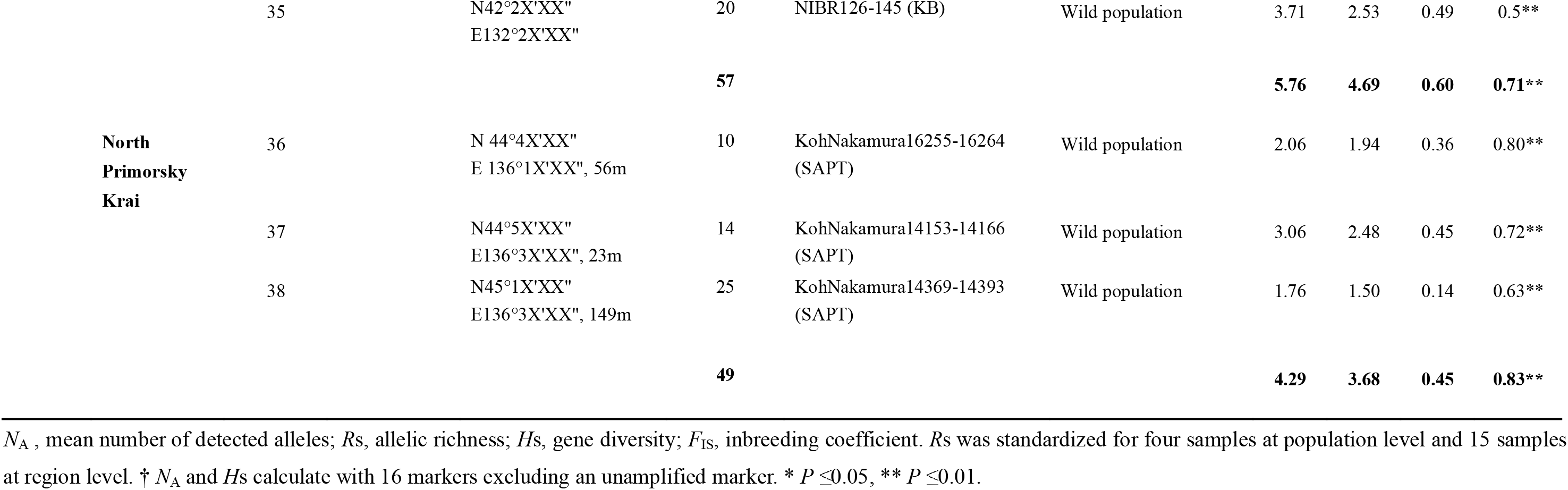
Geographic and genetic characteristics of *Lychnis wilfordii* populations from seven regions of four countries, Japan, Korea, northeastern China and Russian Far East.

There are a few conservation studies of this species in Japan (Tamura *et al.* 2016) and Korea (Jae & Sungwon 2016), the latter focusing on conservation activities in Korea. The former study (Tamura *et al.*, 2016) conducted conservation genetic analysis of Japanese populations of *L. wilfordii* using a limited number of microsatellite markers and revealed that Hokkaido populations (northernmost populations of Japan) are distinct from Nagano populations (southernmost populations); but no samples were used from Korea, northeast China or Russian Far East and species range-wide genetic structure was not elucidated.

In this study we conducted population genetic analyses using 17 nuclear microsatellite (nSSR) markers, and molecular dating using cpDNA sequence data. The aims of this study are 1) to elucidate geographic genetic structure of *L. wilfordii* covering the species range, 2) to test possible scenarios of migration in northeast Asia, and 3) to develop logical plans for effective conservation based on geographic genetic structure and ecological information.

## MATERIALS AND METHODS

### Materials

*Lychnis wilfordii* is a monoecious perennial herb. This is a diploid species (Kruckeberg, 1960) and reproduces sexually with seeds and asexually with rhizomes (Kozhevnikov *et al.*, 2015; Tamura *et al.,* 2016). Although the seeds usually disperse by gravity, they have many hooked spines on the surface (Nakayama *et al.,* 2006) and can be dispersed long-distance by animals.

This species is designated as endangered species in Japan and Korea not only at national but also at prefectural levels: in Japan, Cr (critically endangered), A (most important and rarest wild species), and EN (endangered) in Hokkaido, Aomori, and Nagano prefectures, respectively (Hokkaido prefecture, 2001, 2016; Aomori prefecture, 2010; Nagano prefecture, 2014, 2017). The numbers of individuals/populations are estimated in Hokkaido (339 individuals in 10 populations, author’s unpublished data), Aomori (a few individual in one population; Aomori prefecture, 2010), Nagano (ca. ten populations; Ministry of the environment of Japan, 2015), and Korea (< 300 individuals in three populations; National Institute of Biological Resources, 2014).

Samples were collected covering the species range in northeast Asia, where geographically isolated seven regions are recognized, i.e., Hokkaido, Aomori, Nagano of Japan, Gangwon-do of Korea, Jilin of China, and south Primorsky Krai and north Primorsky Krai of Russian Far East (Table 1). Field collection was conducted in Hokkaido (103 individuals/10 populations), Nagano (55/6), Gangwon-do (65/3), Jilin (98/10), south Primorsky Krai (57/4), and north Primorsky Krai (49/3) (Table 1; Fig 2). In each population, samples were collected > 50 cm apart from each other not to collect the same clones. In addition, for recently extinct populations in Karuizawa, Nagano prefecture of Japan (Hara *et al.,* 1974; Ikeda, 1997), we utilized herbarium specimens (three individuals) and *ex-situ* collection of a botanic garden (two individuals) (Table 1). The only known population in Aomori prefecture of Japan is in a Japanese military base and inaccessible, and a herbarium specimen was used. In total, 433 individuals from 38 populations were studied from the four countries. In cpDNA sequence analysis, a portion of these samples were used. Voucher specimens of our collection were deposited in the herbaria of Botanic Garden, Hokkaido University (SAPT) and National Institute of Biological Resources (KB).

### DNA extraction, nSSR genotyping, cpDNA sequencing

Total genomic DNA was extracted from silica gel-dried leaves, using the CTAB method (Doyle & Doyle, 1987). In the nSSR analysis, DNA samples were genotyped for 17 nSSR loci by using primer pairs developed for *L. wilfordii* (Kim *et al.* 2018). PCR was conducted using forward primers attached M13-tail (5’–GTA AAA CGA CGG CCA GT–3’), M13 primer labeled with the fluorescent dyes 6-FAM, VIC, NED, or PET (Applied Biosystems, Foster City, California, USA), and reverse primers. PCR reactions were performed in 20 µl total volume with the following reagents: ca. 10 ng genomic DNA, 1unit Taq DNA polymerase master mix (Ampliqon, Rødovre, Denmark), 0.3 µl M13-tailed forward primer, 0.6 µl reverse primer, 0.6 µl fluorescent-labeled M13 primer (Table 2) and 2 % dimethyl sulphoxide (DMSO). The PCR cycle conditions were as follows: initial denaturation at 95 °C for 10 min; 35 cycles of denaturation at 95 °C for 30 sec, annealing at Ta °C (Table 2) for 30 sec, and elongation at 72 °C for 30 sec; and final extension at 72 °C for 7 min. For Lw80 marker, 2nd PCR (10 cycles) was conducted using 0.5 µl 1st PCR product (amplified without using fluorescent dye) diluted with sterilized water by three times. The PCR product size was measured with the size standard 600LIZ using an ABI Prism 3130 DNA analyzer (Applied Biosystems, Foster City, CA, USA) and Peak Scanner software v1.0 (Thermo Fisher Scientific, MA, USA).

**Table 2.** Characteristics of 17 microsatellite markers developed for *Lychnis wilfordii* (Kwak *et al.*, in preparation).

| Locus | Repeat motif | Size range | Annealing temperature<br>(Ta, °C) | Fluorescent dye |
| --- | --- | --- | --- | --- |
| Lw06† | AT | 132-199 | 60-55* | 6-FAM |
| Lw07† | GT | 380-394 | 60-55* | NED |
| Lw08† | CA | 409-419 | 55 | PET |
| Lw16 | AG | 207-243 | 55 | 6-FAM |
| Lw21† | AG | 249-280 | 52 | VIC |
| Lw25 | AC | 394-404 | 55 | PET |
| Lw26 | AC | 392-402 | 55 | PET |
| Lw37 | TG | 236-274 | 65-60* | VIC |
| Lw38 | TG | 180-208 | 65-60* | 6-FAM |
| Lw52 | ATC | 218-254 | 55 | VIC |
| Lw57 | ACA | 386-424 | 55 | PET |
| Lw59 | TTC | 329-374 | 55 | NED |
| Lw68† | CT | 245-265 | 50 | NED |
| Lw69 | AG | 200-216 | 55 | 6-FAM |
| Lw80 | AG | 221-239 | 60-55*(1stPCR) /<br>55(2ndPCR) | NED |
| Lw83† | GT | 370-392 | 55 | NED |
| Lw91 | TA | 304-316 | 55 | VIC |

\* Indicates touchdown PCR. The annealing temperature of the first 10 cycles was decreased 0.5 °C per cycle. † Reverse primers attached with PIG-tail sequence (5'-GTTTCTT-3') to promote full adenylation of PCR fragments.

For DNA sequencing, eight cpDNA regions (Table 3) were PCR amplified. PCR reactions were performed in 25 µl total volume with the following reagents: ca. 10 ng genomic DNA, 1unit Taq DNA polymerase master mix (Ampliqon), 1µl each primer and 4 % DMSO. The PCR cycle conditions were 95 °C for 4 min, 30 cycles of 94 °C for 50 sec, Ta °C (Table 3) for 50 sec and 72 °C for 40 sec, and final extension at 72 °C for 10 min. The PCR fragments were purified with isopropanol precipitation. Purified PCR fragments were used as template for cycle sequencing reactions with the same primers used in the PCR, and direct sequencing was performed on an ABI Prism 3130 DNA analyzer.

**Table 3.**
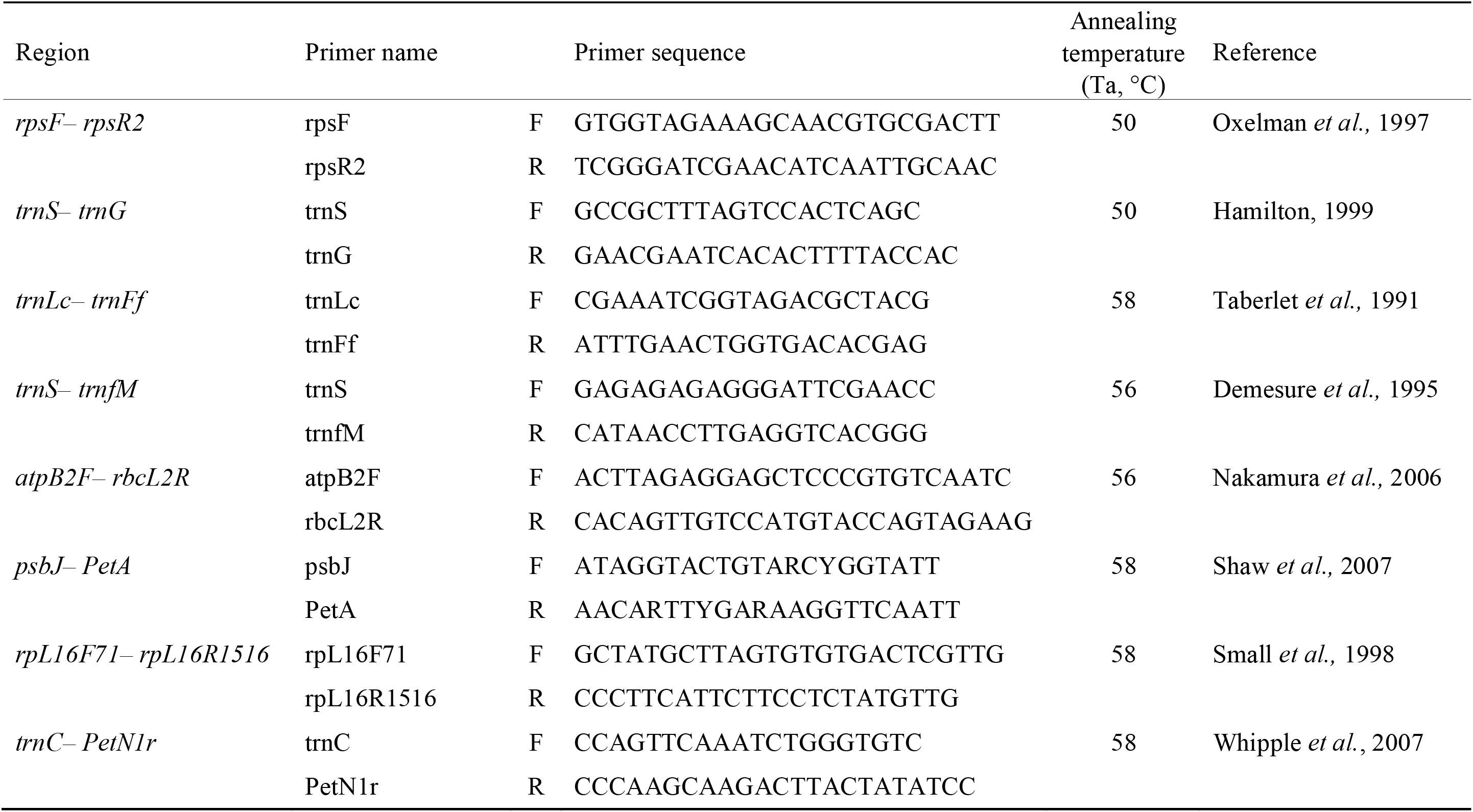
Characteristics of eight cpDNA markers for sequencing.

### Analyses based on nSSR data

#### Genetic profile

The following diversity and inbreeding parameters were computed for populations (n ≥ 4) and each of the six regions (excluding Aomori) using FSTAT 2.9.3.2 (Goudet, 2002): the total number of detected alleles (*N*_A_), allelic richness (*R*s, number of alleles independent of sample size, standardizes for the smallest number of individuals per unit using rarefaction. For populations/regions, it was rarefied to 4/15 samples with 15/16 markers excluding markers unamplified in six populations/one region), genetic diversity (*H*s) and the average inbreeding coefficient (*F*_IS_) across all the loci. For *F*_IS_, the significance of departures from zero was tested with 10 200 randomizations.

#### Population genetic structure

Population genetic structure was analyzed using a Bayesian clustering algorithm implemented in STRUCTURE 2.3.4 (Pritchard *et al.,* 2000, 2010), which probabilistically assigns individuals to clusters based on their multi-locus genotypes by minimizing Hardy-Weinberg and linkage disequilibria. To estimate the most probable number of clusters (*K*), we first conducted a preliminary analysis using about half of the samples (237 samples) from the whole species range, changing *K* from 1 to 20, and the optimal *K* was suggested to be below 10. We therefore set the highest *K* = 10, which is greater than the number of the geographically isolated regions (i.e., Hokkaido, Aomori, Nagano, Gangwon-do, Jilin, south Primorsky Krai, and north Primorsky Krai). The analysis was performed assuming the admixture model and correlated allele frequencies among clusters with default parameter settings. Twenty independent runs for each *K* were carried out to verify the consistency of results, with 2 000 000 MCMC replications after a burn-in length of 500 000 replications. To estimate the most probable *K*, the statistic Δ*K* based on the second-order rate of change of the posterior probability of the data ln *P*(*D*) with respect to *K* (Evanno *et al.*, 2005) was computed as average over 20 runs using STRUCTURE HARVESTER (Earl & vonHoldt, 2012). For the value of *K* selected, the symmetric similarity coefficient (SCC: *H’*) between all pairs of 20 runs, calculated with CLUMPP 1.1.2 (Jakobsson & Rosenberg, 2007), was > 0.45, and membership coefficients (*Q*) of each individual was calculated as average over 20 runs using CLUMPP. The *Q* values were visualized using DISTRUCT (Rosenberg, 2004).

Genetic distance *D*_A_ (Nei *et al.,* 1983) among all 38 populations and among six regions (except Aomori with only one sample) were calculated based on the nSSR data using Populations 1.2.32 (Langella, 2010) and subjected to principal coordinates analysis (PCoA) using GenALEx 6.5 (Peakall & Smouse, 2012).

### Analyses based on cpDNA data

#### MRCA dating

The sequences were aligned using GeneStudio 2.2 (GeneStudio Inc., Suwanee, Georgia) and then adjusted manually using BioEdit v7.2.5 (Hall, 1999). The age of the most recent common ancestor (MRCA) was estimated using BEAST 1.8.4 (Drummond *et al.,* 2012). The appropriate evolutionary model for the cpDNA data was SYM+G, as selected using PAUP* ver. 4.0b10 (Swofford, 2002) and MrModeltest 2.3 (Nylander, 2008), based on the Akaike Information Criterion (AIC). An empirical base frequency and a lognormal relaxed clock rate were used. For molecular dating, a previous study used a split between the subfamilies Alsinoideae and Caryophylloideae as a fossil-calibrated node to estimate the age of the MRCA of the genus *Lychnis*, using a coding region of mat*K* (Gizaw *et al.,* 2016); but a substitution rate for non-coding cpDNA region is not available for this group. We therefore referred to previous reports on non-coding cpDNA substitution rates (per site per year) for other herbaceous plants, most of which have a minimum generation time of 1-3 years (minimum value = 1.30 x 10^-9^, mean value = 4.77 x 10^-9^, maximum value = 8.24 x 10^-9^) (Richardson *et al.,* 2001). To obtain the similar distribution to that of these reported values, a gamma distribution prior was employed for clock rate with an initial value of 4.77 x 10^-9^, a shape value of 4.095, a scale value of 9.2 x 10^-9^ and an offset value of 0.00, which had the median of 3.47 x 10^-9^ and 95 % range of the distribution between 1.05 x 10^-9^ and 8.20 x 10^-9^. A starting tree was generated using random starting tree option. Markov chain Monte Carlo (MCMC) chain length was 200 million steps and parameter values were logged every 1000 steps. The initial 10 % of the sampled parameter values were discarded as burn-in. Convergence of MCMC algorithms and effective sample size (ESS > 500) for each parameter were checked using TRACER 1.6 (Rambaut *et al.,* 2013). A maximum clade credibility tree was estimated with a posterior probability limit (*PP*) of 0.5 by TreeAnnotator ver. 1.8.4 (Drummond *et al.,* 2012), and the estimated age of the MRCA (mean age and 95 % highest posterior density [HPD] interval) was checked on FigTree ver. 1.4.2 (Rambaut, 2014).

## RESULTS

### Analyses based on nSSR data

#### Genetic profile

The genetic diversity estimates at region level indicated comparatively lager values in the continent and peninsula, i.e., north Primorsky Krai (*R*s = 3.68, *H*s = 0.45), south Primorsky Krai (4.69, 0.60), Jilin (5.72, 0.59), and, Gangwon-do (3.72, 0.60) than in regions in Japan, i.e., Hokkaido (1.49, 0.14) and Nagano (1.8, 0.19) (Table 1). *F*_IS_ values ranged from −0.56 to 0.75 at population level and 0.59 to 0.92 at region level (Table 1). *F*_IS_ values were significantly different from zero in most populations (Table 1).

#### Population genetic structure

In the STRUCTURE analysis, *K* = 3 and 5 were supported as the most and second probable numbers of clusters by Δ*K* (Fig. S). When *K* = 3 (Fig. 3), Hokkaido (populations no. 1–10) and Jilin (no. 22–31) almost exclusively belonged to the blue and yellow clusters, respectively. Nagano (no. 12–18) chiefly belonged to the red cluster, but secondarily to the blue cluster. Gangwon-do populations belonged chiefly to the red (no. 19) or yellow (no. 20, 21) clusters and secondarily vice versa. South and north Primorsky Krai populations primarily belonged to the red (no. 32, 35), yellow (no. 33, 34, 36), and blue (no. 37, 38) clusters but also belonged to the other two clusters with varying degrees. When *K* = 5, the genetic difference among the regions became clearer than *K* = 3. In detail, the populations from Nagano (excluding no. 18), Gangwon-do (excluding no. 19), Jilin, and Hokkaido almost entirely belonged to the red, pink, yellow, and blue clusters, respectively. The majority of south Primorsky Krai (no. 33–35), north Primorsky Krai (no. 36–38), and Aomori (no. 11) populations mainly belonged to the green cluster, while partly belonged to the yellow, pink and/or blue clusters. The result showed non-negligible *Q* values for more than two clusters in Nagano (no. 18, mainly to the red but partly to the green and pink), Gangwon-do (no. 19, mainly to the green and pink but partly to the yellow and blue), and south Primorsky Krai (no. 32, mainly to the pink but partly to the green and yellow).

**Figure 3.**
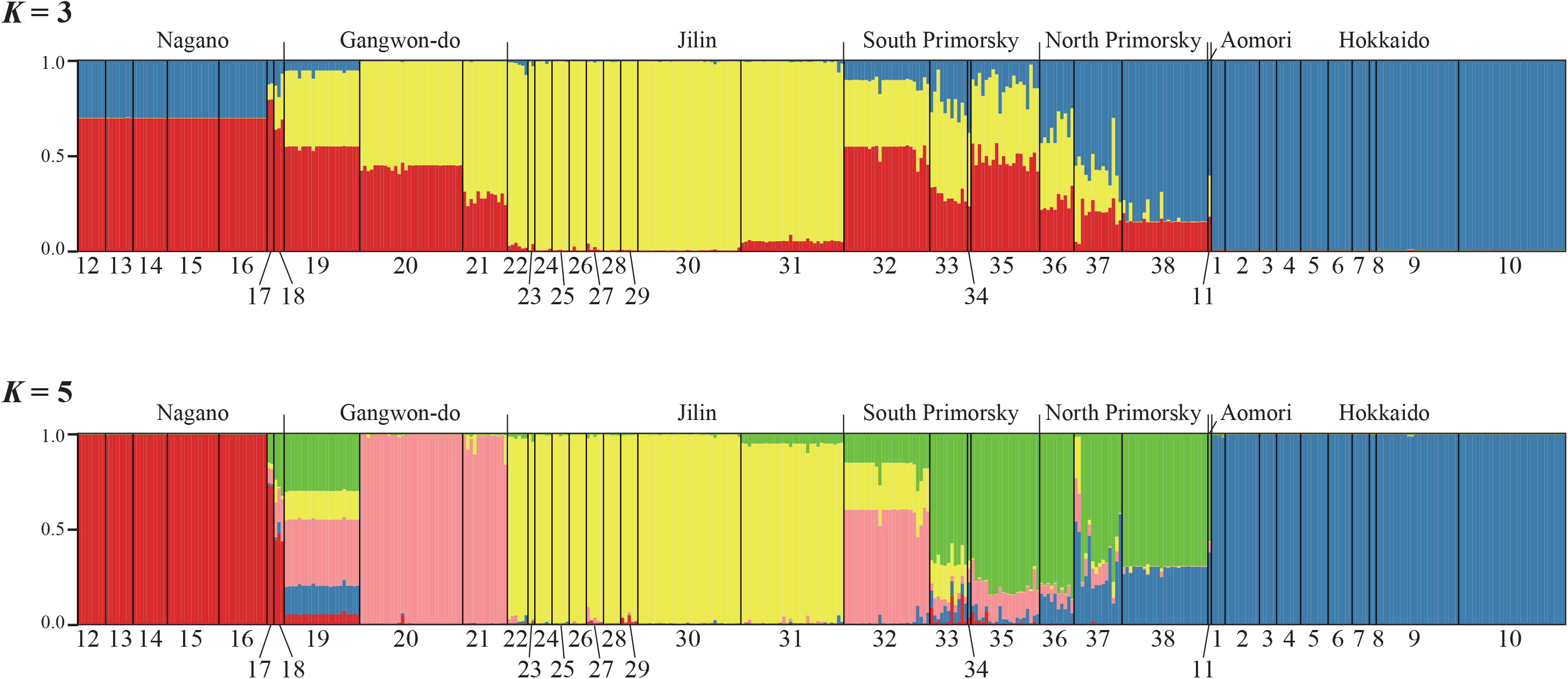
Results of the Bayesian clustering analyses for *Lychnis wilfordii* based on nSSR data. Bar plots show the membership coefficients *Q*. Each individual is represented by a vertical line partitioned into coloured components corresponding to its mean membership coefficients over 20 runs of the STRUCTURE analysis. For the population numbers (1-38), see Table 1.

*D*_A_ genetic distance among populations ranged from 0 to 1.00, with the mean of 0.67 (Table S). The value was large comparatively (here defined as ≥ 0.95) between Nagano (no. 17) and Jilin (no. 22, 24, 26, 28 and 30) (*D*_A_ = 0.95-1.00), and Gangwon-do (no. 19) and Jilin (no. 22, 23, 26 and 27) (*D*_A_ = 0.96-0.97), on the other hand the value was small comparatively (here defined as ≤ 0.05) between populations in Hokkaido (population no. 1 and 5, *D*_A_ = 0.05; 2 and 4, 0.00; 2 and 5, 4 and 5, 0.01), and in Nagano (no. 14 and 15, 0.00). The PCoA plots based on *D*_A_ is shown (Fig. 4). The first and second axes extracted 28.39 % and 27.43 % of the total genetic variation. Plots of populations clustered roughly corresponded to the five groups, i.e., Nagano (no. 12-17), Gangwon-do (no. 19-21), Jilin (no. 22-31), south and north Primorsky Krai (no. 32-38), and Hokkaido plus Aomori (no. 1-11); note that one Nagano population (no. 18) was exceptional and plotted closer to Gangwon-do populations. The plots indicated genetic proximity between Nagano and Gangwon-do populations, between south and north Primorsky Krai, and Hokkaido plus Aomori populations, and among Jilin, Gangwon-do, and south and north Primorsky Krai populations, whereas Hokkaido and Nagano populations were plotted distantly. *D*_A_ genetic distance among the six regions (excluding Aomori with only one sample; Table 4) was comparatively large (here defined as > 0.70) between Hokkaido and Gangwon-do (*D*_A_ = 0.83), Nagano and Jilin (0.81), Nagano and north Primorsky Krai (0.74), and Hokkaido and Nagano (0.70); whereas it was comparatively small (< 0.40) between Hokkaido and north Primorsky Krai (0.34) and north Primorsky Krai and south Primorsky Krai (0.39). The PCoA plots based on *D*_A_ for these six regions is shown (Fig. 5). The first and second axes extracted 40.69 % and 35.19 % of the total genetic variation. The plots showed the genetic proximity among Jilin, south Primorsky Krai, and north Primorsky Krai populations, and between north Primorsky Krai and Hokkaido populations. Hokkaido and Nagano were again plotted distantly.

**Figure 4.**
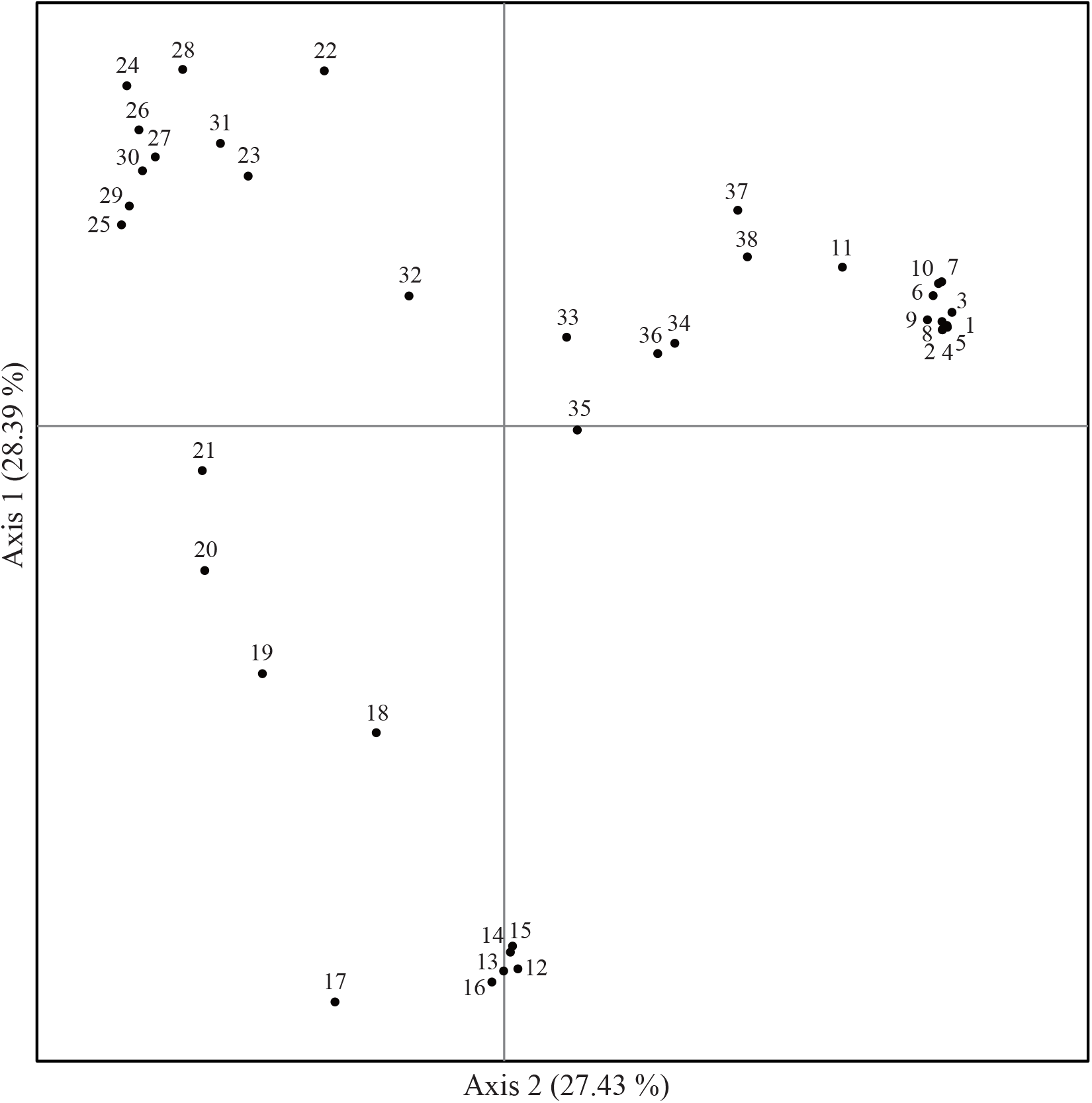
Plots of principal coordinates analysis (PCoA) of *Lychnis wilfordii* using the nSSR data, based on *D*_A_ genetic distance among populations. Populations 1-10 are from Hokkaido; 11 is from Aomori; 12-18 are from Nagano; 19-21 from Gangwon-do; 22-31 from Jilin; 32-35 from south Primorsky; 36-38 from north Primorsky. For details of the populations, see Table 1.

**Figure 5.**
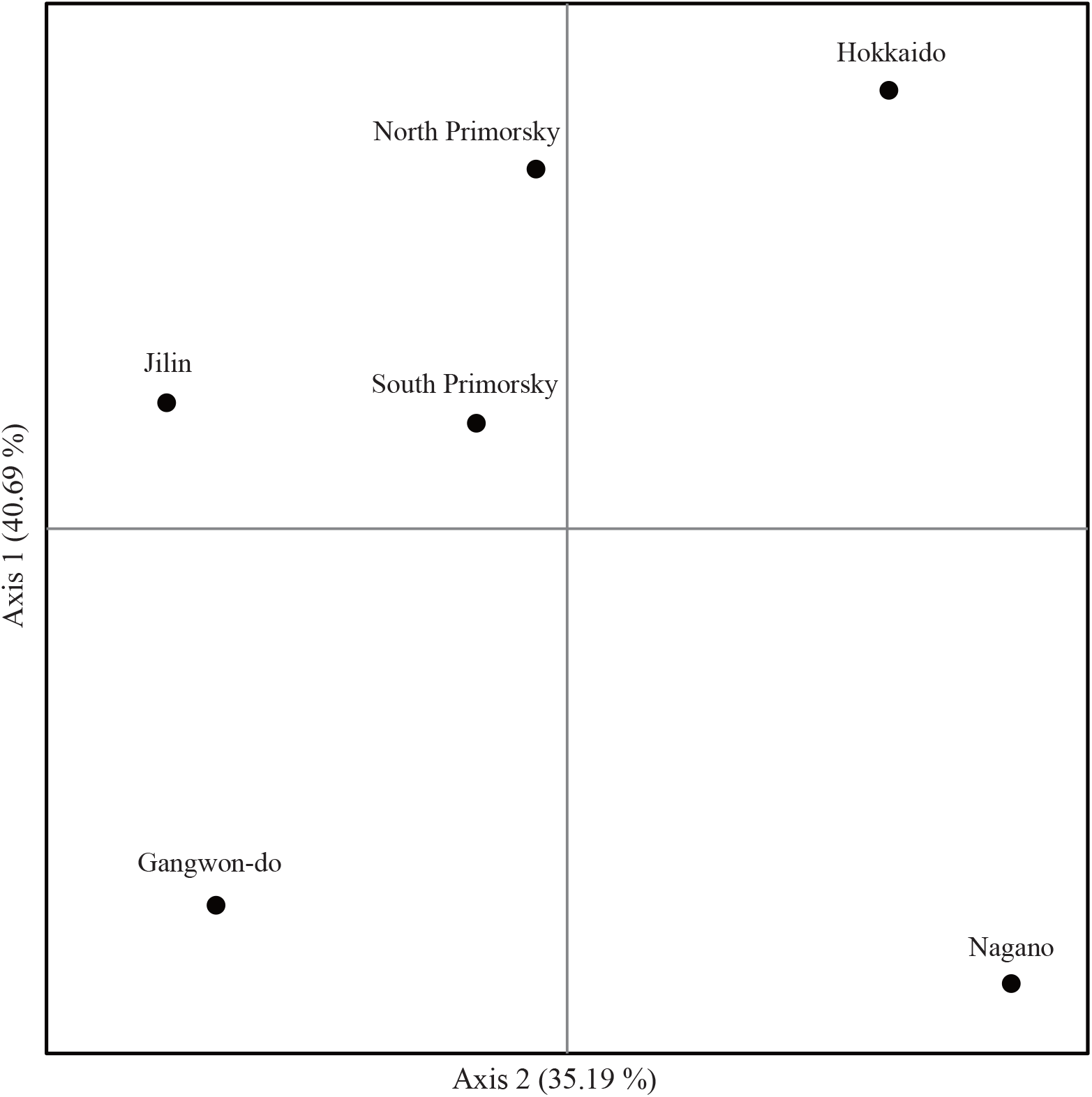
Plots of principal coordinates analysis (PCoA) of *Lychnis wilfordii* using the nSSR data, based on *D*_A_ genetic distanve among six regions. Aomori region was excluded due to n = 1. For details of the regions, see Table 1.

**Table 4.** Pairwise genetic distance *D*_A_ among six regions.

| Region | Hokkaido | Nagano | Gangwon-do | Jilin | South Primorsky<br>Krai | North Primorsky<br>Krai |
| --- | --- | --- | --- | --- | --- | --- |
| Hokkaido | 0.00 |  |  |  |  |  |
| Nagano | 0.70 | 0.00 |  |  |  |  |
| Gangwon-do | 0.83 | 0.67 | 0.00 |  |  |  |
| Jilin | 0.67 | 0.81 | 0.58 | 0.00 |  |  |
| South Primorsky<br>Krai | 0.54 | 0.64 | 0.54 | 0.51 | 0.00 |  |
| North Primorsky<br>Krai | 0.34 | 0.74 | 0.62 | 0.53 | 0.39 | 0.00 |

#### MRCA dating based on cpDNA

The aligned length of cpDNA data was 4693 bp, where only one single nucleotide difference was found between Chinese samples and the other ingroups. Thereby, the data provided little resolution within the ingroup clade (*PP* = 1). The Bayesian molecular dating based on cpDNA estimated the mean age of the MRCA of *L. wilfordii* as 0.85 million years (Myr), i.e., the base of the Late Pleistocene, and 95 % highest posterior density (HPD) interval as 0.005–0.42 Myr, i.e., the middle Pleistocene to the Holocene.

## DISCUSSION

### Geographic genetic structure

The population genetic analyses using 17 nSSR markers were conducted to elucidate geographic genetic structure of *L. wilfordii* covering the whole species range in northeast Asia. Based on the results of the STRUCTURE (*K* = 5) and PCoA analyses, five genetically distinct groups were recognized, namely, Nagano, Gangwon-do, Jilin, north and south Primorsky Krai plus Aomori, and Hokkaido (Figs. 3, 4). In the PCoA (Fig. 4), Hokkaido populations are clustered in the proximity to Aomori and north Primorsky Krai populations of the continent but away from Nagano populations of the same Japanese Archipelago. On the other hand, Nagano populations are plotted close to Gangwon-do populations of the Korean peninsula, and Gangwon-do populations are to south Primorsky Krai populations. These close relationships also represented at STRUCTURE (Fig. 3), which was indicated non-negligible *Q* values for more than two clusters at population no. 18 in Nagano, 19 in Gangwon-do, and 32 in south Primorsky Krai. PCoA plots of Jilin populations are close to those of south Primorsky Krai populations. The PCoA plots of the five genetic groups roughly correspond to their geographic distribution in northeast Asia (Figs. 2, 4). This is more evident in the PCoA at the regional level (Fig. 5). This suggests that the geographic genetic structure was formed reflecting migratory history of the species, as discussed later.

It is contrasting that in the continent and the peninsula, populations of the same country (i.e., Korea, northeast China, or Russian Far East) clustered together and formed the genetically distinct group, whereas in the archipelago, the Japanese populations did not cluster together but highly differentiated between Hokkaido plus Aomori and Nagano (Figs. 4, 5). It is worth to note that the genetic distance between Hokkaido and Nagano (*D*_A_ = 0.70, Table 4) was larger than between Hokkaido and north Primorsky Krai (0.34) and between Nagano and Gangwon-do (0.67) crossing national borders and the natural barrier of sea. Although the genetic difference between Hokkaido and Nagano was recognized in the previous study (Tamura *et al.*, 2016), the present study covering the whole species range revealed that the genetic difference between Hokkaido and Nagano are relatively large among regions of the other countries. What the geographic genetic structure revealed here suggests for the conservation of *L. wilfordii* is discussed below.

### Possible scenario of migration

To discuss the migratory history of *L. wilfordii* in northeast Asia, its ancestral area should be elucidated. The nSSR analysis indicated that the genetically recognized groups in the continent and peninsula parts (north and south Primorsky Krai, Jilin, and Gangwon-do) had higher genetic diversity compared to those in the Japanese Archipelago (Hokkaido and Nagano) (Table 1). In general, progenitor populations in ancestral areas have higher genetic diversity than derivative populations due to founder effect in the latter (Frankham, 1996; Hewitt, 1996; Ibrahim *et al.,* 1996), although this pattern can be disrupted by other population genetic processes such as bottleneck events in ancestral populations, gene flow/recurrent migration to derivative populations, and natural selection (e.g. Hiramatsu *et al.,* 2001; Tremetsberger *et al.,* 2003; Morrell *et al.,* 2003). Additionally, genus *Lychnis* is distributed widely from Europe, via Africa, to northeast Asia (Lu *et al.,* 2001), and the genus have likely expanded its range from the Eurasian continent to Africa (Popp *et al.,* 2008; Gizaw *et al.,* 2016), suggesting that Eurasian species are relatively old in the genus. *Lychnis wilfordii* has the phylogenetically closest species in the Asian mainland (Ullbors, 2008). Based on these data and the circumstantial evidence, it is highly likely that the Asian continent is the ancestral area of *L. wilfordii* and the species expanded to the Japanese Archipelago. As discussed above, in the PCoA based on *D*_A_ distance (Figs. 4, 5), the plots of Nagano, Gangwon-do, Jilin, north Primorsky Krai, south Primorsky Krai, Aomori, and Hokkaido roughly corresponded to their geographic distribution in northeast Asia (Fig. 2), suggesting that the geographic genetic structure was formed reflecting migratory history of the species. Considering that Hokkaido and Aomori populations are genetically close to north Primorsky Krai populations and that Nagano populations are to Korean populations, *L. wilfordii* likely migrated from the Asian continent to the Japanese Archipelago using two routes: north route from Russian Far East to Hokkaido and Aomori, and south route from the Korean Peninsula to Nagano.

The age of the MRCA of *L. wilfordii* was estimated as 0.085 Myr (0.005–0.42 Myr) and this corresponds to the middle Pleistocene–the Holocene. During the Last Glacial Maximum (LGM), the climate was cooler and the sea level was ca. 85-130 m below the present level, and the huge ice sheets developed in high-latitude areas of the Northern Hemisphere and Antarctic (Tsukada, 1983; Oba & Irino, 2012). It was suggested that there was landbridge formation between the Korean Peninsula and southwestern Japan during the Pleistocene including the complete closure of the strait and the inundated landbridge (Oba *et al.,* 1991; Millien-Parra & Jaeger, 1999; Kim *et al.,* 2000; Lee & Nam, 2003; Lee *et al.,* 2008; Oba & Irino, 2012). Many studies suggested that various plant lineages migrated using the landbridge between the Korean Peninsula and southwestern Japan (e.g. Aizawa *et al.*, 2007, 2009; Kikuchi *et al.,* 2010, 2013; Qi *et al.,* 2012). It is highly likely that the south-route migration of *L. wilfordii* from Korea to Japan was facilitated by that landbridge, even if it was split by a narrow strait. On the other hand, concerning the north route, there was no direct land connection between the continental part of Russian Far East and Hokkaido, although Sakhalin Island connected them, during the Pleistocene (Minato & Ijiri, 1984, Yasuda, 1984; Oba *et al.*, 1991; Oba & Irino, 2012). However currently *L. wilfordii* is not distributed in Sakhalin (Institute of biology and soil science, Russian academy of sciences, Far Eastern branch, 1996), and it would not be parsimonious or rational to postulate its extinction in Sakhalin because *L. fulgens*, that sometimes grows sympatrically with *L. wilfordii* in Primorsky Krai (Institute of biology and soil science, Russian academy of sciences, Far Eastern branch, 1996; author’s observation), is found in Sakhalin (Sugawara, 1975). This means that *L. wilfordii* likely expanded the distribution via the north route by long-distance dispersal. Bird dispersal is a highly probable scenario considering the seed morphology having many hooked spines on the surface (Nakayama *et al.*, 2006). Bird dispersal may also have happened between Gangwon-do (no. 19) and south Primorsky Krai (no. 32, 33 and 34), for which genetic affinity was indicated by the STRUCTURE and PCoA analyses despited the large geographic distance.

The north and south migratory routes has long been recognized to have played a fundamental role in shaping the flora of the Japanese Archipelago (Good, 1974; Maekawa, 1974; Takhtajan, 1986). The migration of *L. wilfordii* via both the north and south routes, however, is a rare example of two-way migration by a single species. Previously, this was revaled by molecular data only for a northeast Asian spruce *Picea jezoensis* (Aizawa *et al.*, 2007, 2009), although it was hypothesized, but without verification, in several plant species (Lihová *et al.*, 2010; Kikuchi *et al.*, 2010, 2013). The connection between the circular landform of Japan-Korea-northeast China-Russian Far East region and the potential two-way migration of a species is an interesting issue for further study in conservation biology as well as biogeography in northeast Asia.

### Logical plans for effective conservation

In conservation biology, conservation units (evolutionary significant units, ESUs, management units, MUs) are defined for effective conservation practice. Based on the geographic genetic structure, it became clear that there are five genetically distinct groups in *L. wilfordii*, i.e., Hokkaido, Nagano, Korea, northeast China, and Russian Far East. In particular, Hokkaido and Nagano populations should be distinguished as different ESUs because Hokkaido and Nagano populations are at the different ends of the two migratory routes based on the migration scenario, although these two regions belong to the same country. Outbreeding depression can happen as a result of gene flow between diverged populations, especially when the species have limited dispersal ability and populations are adapted to local environments (Frankham *et al.*, 2002). The dispersal of this species between the regions (i.e., Hokkaido, Nagano, Korea, northeast China and Russian Far East) is limited. Its habitats are wetlands and environments are similar among populations across the species range (Shimizu, 1997; Lu *et al.*, 2001; National Institute of Biological Resources, 2014; Tamura *et al.,* 2016), however, climatic conditions, specifically temperature in winter and day length, are different among the regions at different latitudes. Therefore the crossing between ESUs can potentially cause outbreeding depression. It is worth to note that there found a slight morphological difference between Hokkaido and Nagano populations; the leaf edge is flat in the former but waved in the latter. More detailed morphological studies, together with cross-fertility tests of the genetic groups recognized in the present study, are vitally needed to further improve conservation planning.

*Lychnis wilfordii* is the target of conservation activities in Japan and Korea, but in northeast China and Russian Far East, no special conservation measures are currently taken for *L. wilfordii* because this species is not designated as an endangered species in China or Russia. This study revealed the five genetically distinct groups and defined the ESUs of Hokkaido and Nagano. Considering that the continental populations are potentially migratory source for the island populations in an evolutionary time scale, transnational conservation of *L. wilfordii* in northeast Asia is vitally needed.

---

**Figure S.**
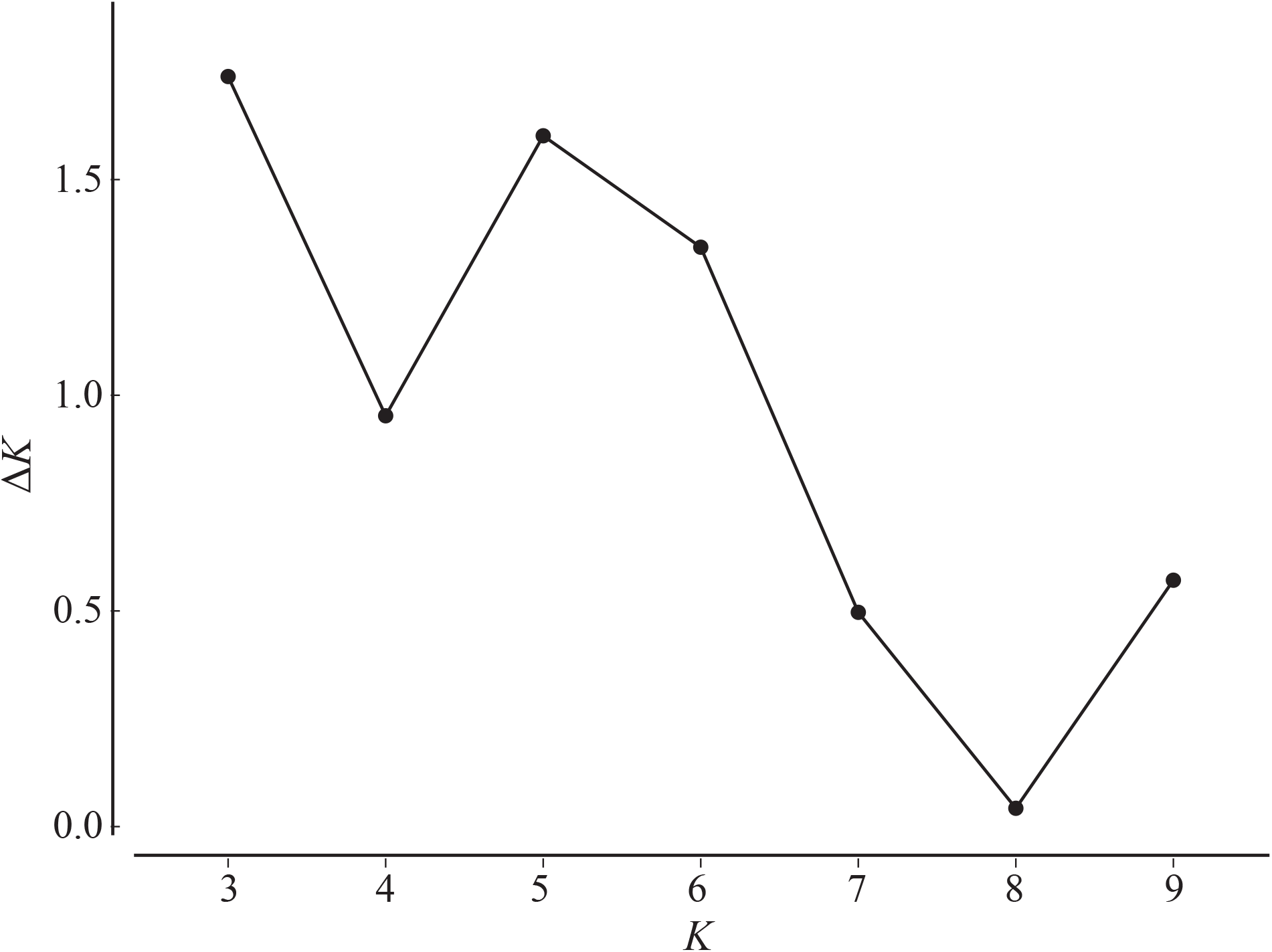
Results of the Bayesian clustering analyses of *Lychnis wilfordii* based on the nSSR data. Plot of the statistic Δ*K* (*K*, number of clusters) based on the second-order rate of change of ln *P*(*D*) with respect to *K*. Bar plots showing the membership coefficients *Q* at *K* = 3 and 6 are provided in Fig. 3.

